# GAE-Δ: A Graph-Learning Framework for Gene Network Rewiring and Clinical Outcome Prediction from Multi-Omics Data

**DOI:** 10.64898/2026.05.21.726880

**Authors:** Zhiyong Tang, Zhe Chen, Mengting Chen, Yihua Wang, Sarah Ennis, Mahesan Niranjan, Rob M. Ewing

## Abstract

Cancer progression and outcomes are driven in part by changes to molecular networks that result from genetic and/or environmental perturbations. These network changes manifest across multiple interconnected network layers and include accumulation of somatic mutations, altered proteinprotein interactions and dysregulated gene-expression. Here we describe a graph autoencoder–based framework (Graph Autoencoder-Delta (GAE-Δ)), for characterizing phenotype-specific gene role shifts across multi-omics data. Given samples stratified into two contrasting phenotypic groups and a prior gene interaction network, GAE-Δ constructs group-specific gene graphs for each omics modality and trains, for each modality, a single graph autoencoder jointly on both group graphs, so that the two group-conditional embeddings share a common latent space. Contrasting these embeddings defines a multi-omics embedding-shift representation for each gene that reflects how its network role reorganizes across phenotypic contexts. These gene-level shifts are subsequently used for unsupervised gene prioritization, multi-omics late fusion and sample-level classification. Applied to five TCGA cancer types with a survival endpoint, GAE-Δ achieves competitive or superior predictive performance compared with classical network-based methods and multi-omics matrix-factorisation methods (MOFA+, iNMF), with statistically significant AUC gains over MOFA+ in three of five cohorts and statistical ties on the remaining two. Beyond predictive performance, the consensus shift genes are significantly enriched for known cancer drivers in three of five cohorts (hypergeometric *p* < 0.01; 11–17× fold-enrichment), whereas matrix-factorisation baselines reach *p* < 0.05 in zero of five cohorts (best per-cancer *p* = 0.06), indicating that GAE-Δ captures biological signal that linear factor methods miss. In summary, the GAE-Δ approach provides for both improved outcome classification as well as for biological and mechanistic discovery through deep network-based integration of disease-associated multi-omics data.

## Introduction

Cancer progression involves coordinated dysregulation across multiple molecular layers, including transcriptional programs, epigenetic remodelling and genomic alterations (Hoadley et al., 2018; Sanchez-Vega et al., 2018). Large-scale multi-omics resources such as The Cancer Genome Atlas (TCGA) have enabled systematic dissection of tumour heterogeneity and clinical outcome (Weinstein et al., 2013), yet many prognostic studies treat genes as independent features and primarily capture changes in abundance (van ‘t Veer et al., 2002).

To better exploit systems-level information, a growing body of work has leveraged multi-omics integration and deep learning for cancer prognosis. Unsupervised and supervised autoencoder models have been used to compress high-dimensional multi-omics data into low-dimensional patient embeddings for survival prediction (Chaudhary et al., 2018; Zhang et al., 2018). In parallel, graphbased methods have incorporated prior biological networks—such as protein–protein or functional interaction (FI) networks—using graph convolutional networks (GCNs) to integrate omics features defined on gene graphs (Dai et al., 2025; Li et al., 2022). These approaches improve robustness by explicitly encoding gene–gene relationships, yet they typically construct a single global graph per cohort and use static node or patient embeddings, without quantifying how network organization differs between clinical outcome groups. However, cancer progression is tightly linked to the reorganization of molecular interaction networks, where genes may change their functional context without exhibiting strong changes in bulk expression (Arshad and McDonald, 2021). Quantifying such reorganization requires comparing a gene’s embedding between two phenotype-specific graphs; a fundamental challenge is that embeddings learned from separately trained models reside on different latent manifolds shaped by stochastic initialization and training dynamics (Bronstein et al., 2017), so that naïvely differencing them conflates genuine biological reorganization with non-biological training artifacts, analogous to the domain shift problem in transfer learning. A representation that places both group-specific embeddings in a common latent space by construction is therefore needed to isolate phenotype-specific signals. Three key limitations persist. First, most methods target patient-level embeddings and do not provide gene-level measures of how network roles change between outcome groups, limiting interpretability (Chaudhary et al., 2018). Second, graph construction is typically coupled to a single omics layer, making it difficult to disentangle transcriptional, epigenetic and copy-number contributions (Li et al., 2022). Third, multi-omics fusion is performed at the raw feature or model level rather than at the level of network role changes (Shen et al., 2018). Existing network-based gene prioritization methods—including G2Vec, NCPR, WuStein and WGCNA—learn embeddings on a single global network shared across all phenotypic groups (Choi et al., 2018). The resulting representations are inherently static topological coordinates: they capture a gene’s structural position within one fixed interaction context, implicitly assuming that molecular interactions remain invariant across clinical states. Yet when network reorganization is the primary signal of interest, what matters is not a gene’s position in any one graph but the displacement between its positions in two contrasting graphs. Genes that undergo substantial functional context changes—shifting from one neighbourhood module to another—may exhibit modest changes in abundance yet large shifts in embedding space. This motivates a transition from static node embeddings to differential embedding shifts as the primary representation for gene prioritization.

A complementary line of work uses multi-omics matrix factorisation methods such as MOFA+ (Argelaguet et al., 2020) and joint NMF variants including iNMF / LIGER (Welch et al., 2019) to recover shared and group-specific latent factors across modalities. These methods offer a principled, convex formulation of multi-group multimodality integration, but they operate at the level of sample-level factor activities rather than gene-level network reorganization, and they assume linear factor structure on raw omics features rather than non-linear reorganization of gene neighbourhoods. We benchmark against MOFA+ and iNMF as the canonical members of this family and demonstrate that GAE-Δ provides complementary — and in three of five cohorts statistically superior — predictive signal, together with gene-level network-rewiring interpretability that factor-loading views structurally cannot produce.

We propose GAE-Δ (Graph Autoencoder-Delta), a general framework that characterizes phenotype-specific gene role shifts in multi-omics data. Given samples stratified into two contrasting phenotypic groups (e.g., good vs poor clinical outcome) and a prior gene interaction network (Milacic et al., 2024), GAE-Δ constructs group-specific gene graphs for each omics modality and trains, per modality, a single graph autoencoder jointly on both group graphs (Kipf and Welling, 2016), so that the two group-conditional embeddings inhabit a common latent space and can be differenced directly. Contrasting these embeddings defines a multiomics embedding-shift representation reflecting how each gene’s network role reorganizes across phenotypic contexts. These shifts are used for unsupervised gene prioritization, multi-omics late fusion and sample-level classification.

Our contributions are fourfold: (i) a phenotype-specific graph autoencoder framework, with a shared-parameter encoder jointly trained across phenotype groups, yielding gene-level measures of network role reorganization applicable to any binary stratification of multi-omics cohorts; (ii) systematic decoupling and late fusion of multiple omics layers in the embedding-shift space; (iii) benchmarking against classical network-based methods and the multi-omics matrix-factorisation family (MOFA+, iNMF) across five TCGA cancer types, with a hypergeometric driver-gene enrichment analysis showing that GAE-Δ’s consensus genes recover known cancer drivers that factor-based baselines miss; and (iv) a TCGA-LIHC case study showing enrichment for well-known oncogenic pathways and interpretable network rewiring.

## Methods

An overview of the GAE-Δ framework is shown in Figure 1.

**Figure 1.**
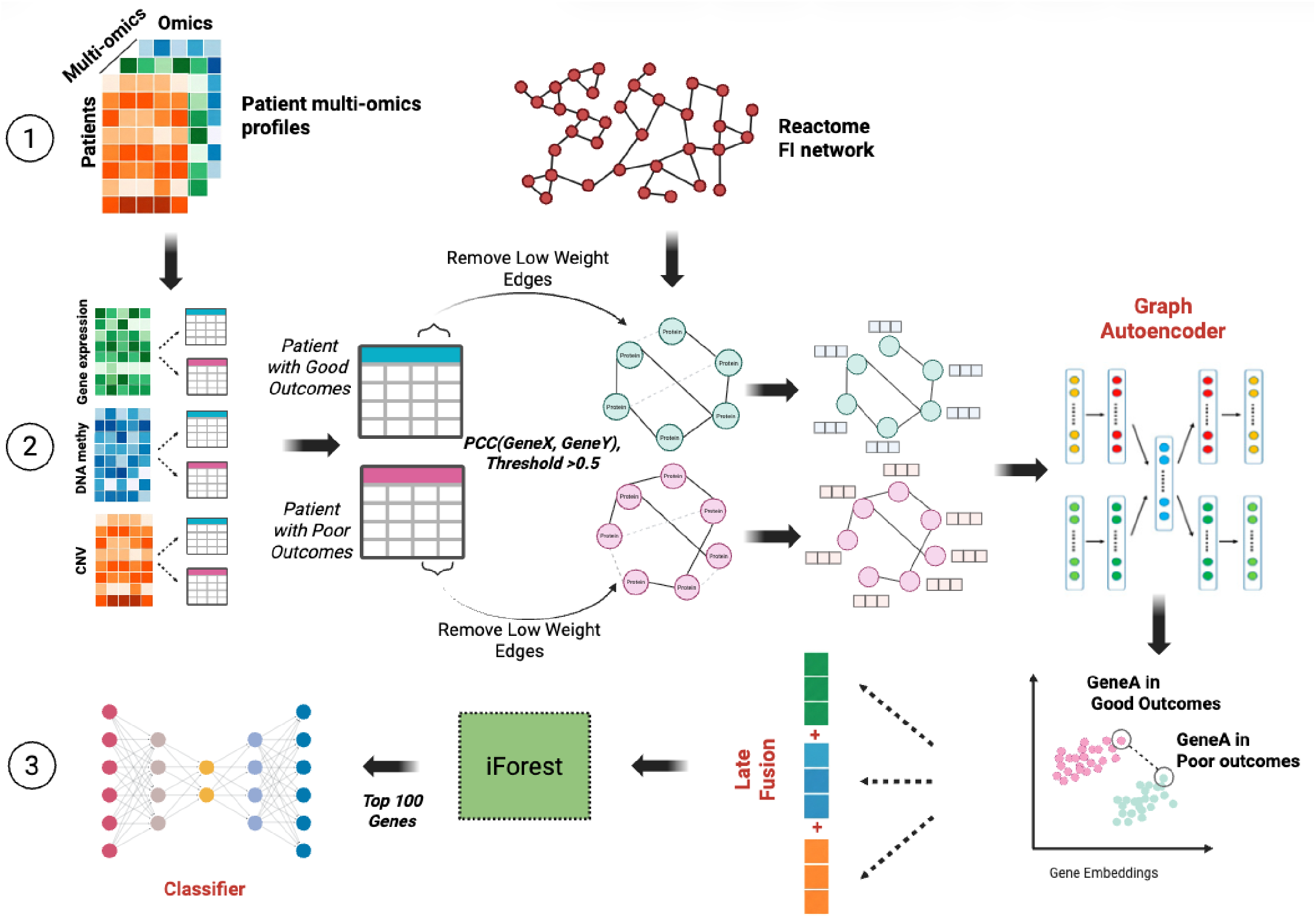
Overview of the GAE-Δ framework. Multi-omics data from samples in two contrasting phenotypic groups (illustrated here with clinical outcome) and a prior gene interaction network are used as input. For each modality and group, phenotype-specific gene graphs are constructed by retaining prior-network edges with high within-group correlation. A single graph autoencoder with shared parameters is trained jointly on both group-specific graphs, producing group-conditional gene embeddings that live in a common latent space by construction. Embedding shifts are computed per gene, concatenated across modalities, and used both for unsupervised gene selection via Isolation Forest and for constructing sample-level embeddings. A multilayer perceptron (MLP) classifier is finally trained on sample embeddings to predict group membership.

### The GAE-Δ framework

The GAE-Δ framework takes as input: (i) a prior gene interaction network *G*_prior_ = (V, E_prior_) encoding known functional relationships among |V| genes; (ii) multi-omics observation matrices *X*^(*c,m*)^ ∈ 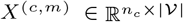for each contrasting phenotypic group *c* ∈ {1, 2} and omics modality *m*; and (iii) a binary group label per sample. The pipeline proceeds in three steps (Figure 1).

#### Step ① : Phenotype-specific graph construction

For each modality *m* and group *c*, an edge (*i, j*) from the prior network is retained if and only if 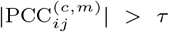, where 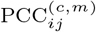 is the Pearson correlation computed from samples in group *c* using modality *m*. This yields a group- and modality-specific graph *G*^(*c,m*)^ that captures co-regulation patterns unique to each phenotypic context. The threshold *τ* is selected via grid search to balance graph connectivity and downstream classification performance.

#### Step ② : Graph autoencoder embedding

Each node is characterized by a 4-dimensional feature vector: (i) within-group mean of standardized omics values, capturing the baseline molecular abundance associated with each group; (ii) within-group standard deviation, reflecting molecular heterogeneity; (iii) node degree in the group-specific graph; and (iv) log-transformed degree log(1+degree), included to stabilize the feature distribution and prevent hub genes from dominating the latent representation. All four features are standardized across nodes within each graph. A single two-layer GCN encoder with shared parameters across phenotype groups is trained *jointly* on both group-specific graphs *G*^(1,*m*)^ and *G*^(2,*m*)^, producing group-conditional readouts from the same latent manifold. With symmetrically normalized adjacency *Ã*^(*c*)^ = (*D*^(*c*)^)^*−*1*/*2^(*A*^(*c*)^+*I*)(*D*^(*c*)^)^*−*1*/*2^, the encoder computes for each group *c*

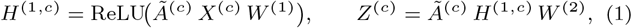

where *W* ^(1)^ ∈ ℝ^*F ×*32^, *W* ^(2)^ ∈ ℝ^32*×*16^ are shared across *c* ∈ {1, 2} (dropout 0.3 on *H*^(1,*c*)^). The decoder reconstructs each group’s edges as 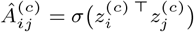. The encoder is trained with the sum of pergroup binary cross-entropy losses, Adam (lr = 10^*−*3^, weight decay 10^*−*4^), and early stopping (patience 30, max 300 epochs). Because both group embeddings are produced by the same parameter set, they live in a common latent manifold by construction, eliminating the inter-run alignment problem that arises when group-specific GAEs are trained independently.

#### Step ③ : Embedding shift

For each gene *g*, we obtain its latent embedding from the group-1 readout 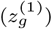 and from the group-2 readout 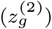. Embeddings are L2-normalized (*z* ← *z/*∥*z*∥_2_) and the raw shift vector for gene *g* is defined as

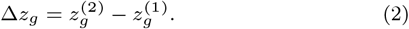

For each modality, this yields a *d*-dimensional shift vector per gene. Shifts from *M* modalities are concatenated into a *Md*-dimensional multi-omics shift vector *s*_*g*_ per gene.

### Optional: KNN residual correction for the independent-encoder variant

As an alternative architectural choice, we also implement an independent-encoder variant in which one GAE is trained per group; here the raw shift Δ*z*_*g*_ may contain a smooth globally shared component reflecting independent-training artifacts. We model this decomposition as Δ*z*_*g*_ = *f* (*g*) + *ε*_*g*_, where *f* (*g*) is a smooth function over the shift space capturing global drift and *ε*_*g*_ is the atypical, gene-specific deviation of interest. To estimate and remove *f* (*g*), K-nearest-neighbour (KNN) regression is applied directly in the raw shift space: for each gene *g*, the *K* nearest neighbours 𝒩_*K*_ (*g*) are identified by Euclidean distance and the local average 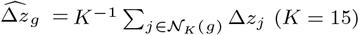 is subtracted to yield the residual *ε*_*g*_.This variant is provided as a switchable option for users who prioritise cross-fold selection-set stability over architectural simplicity; under the shared-encoder default, KNN correction is unnecessary and does not improve predictive AUC (Supplementary Table S11).

### Gene selection via Isolation Forest

To identify genes with pronounced and atypical phenotype-specific role shifts, we applied Isolation Forest to the multi-omics shift representations {*s*_*g*_}. Before model fitting, each dimension of *s*_*g*_ was standardized across genes to equalize contributions from different modalities. Isolation Forest was run with 500 trees and a subsample size of 256. Genes were ranked by anomaly score, and the top *k* genes were selected as candidate phenotype-associated genes for downstream sample-level modelling. This unsupervised procedure prioritizes genes whose shift patterns deviate strongly from the global background, enriching for network roles that reorganize specifically with the contrasting phenotype.

### Sample-level embedding and classification

For the top *k* selected genes 𝒢, each shift vector decomposes as 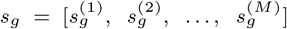 (*d*-dimensional per modality). A sample embedding *u*_*p*_ ∈ ℝ^*Md*^ is constructed by weighting shifts by standardized omics abundances:

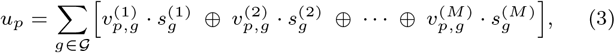

where ⊕ denotes vector concatenation and scalar multiplication is performed elementwise, broadcasted to the embedding dimension. This can be interpreted as representing each sample as a weighted combination of phenotype-specific gene role shifts, linking individual molecular profiles to the network reorganization captured by GAE-Δ.

Binary classification is performed using a two-layer MLP (*Md*→64→1) with ReLU activation and dropout (rate 0.3) after the hidden layer, followed by sigmoid output. The network is trained with binary cross-entropy loss using the Adam optimizer (lr=10^*−*3^, weight decay 10^*−*4^), batch size 32, a maximum of 200 epochs and early stopping with patience 20 based on validation loss.

### Data, baselines and experimental setup

We evaluated GAE-Δ on five cancer types from The Cancer Genome Atlas (TCGA): BLCA, BRCA, CESC, LAML and LIHC (Table 1). Patients were split into good- and poor-outcome groups (*c*=1 and *c*=2, respectively) using median overall survival (OS) as threshold (Katzman et al., 2018). In this application, the prior network *G*_prior_ is the Reactome FI network (Milacic et al., 2024) (14,011 genes, ∼254k interactions), the correlation threshold was set to *τ* = 0.5 (selected via grid search over {0.3, 0.4, 0.5, 0.6}), and *k*=100 genes were selected. Network statistics are reported in Supplementary Table S1. For all predictive modelling, outcome labels were generated within each cross-validation fold using training-set median OS only, ensuring no test-data leakage. Unless otherwise specified, all normalization and feature filtering steps were performed within each fold using only training samples.

**Table 1.** Summary of TCGA cohorts used for evaluation.

| Cancer | $n$ | Good | Poor | Med. OS (d) | Omics genes* |
| --- | --- | --- | --- | --- | --- |
| LIHC | 369 | 188 | 181 | 599 | 14 825 |
| BLCA | 409 | 204 | 205 | 535 | 15 390 |
| BRCA | 1041 | 522 | 519 | 860 | 15 830 |
| CESC | 255 | 128 | 127 | 659 | 15 510 |
| LAML | 200 | 106 | 94 | 365 | 14 280 |

\* Genes with complete multi-omics data.

#### RNA expression

Gene-level RNA-seq profiles were downloaded as HTSeq-FPKM values from TCGA, transformed as log_2_(*x* + 1) and standardized to z-scores per fold.

#### DNA methylation

DNA methylation data were obtained from Illumina HumanMethylation450 arrays as beta values. CpG probes were mapped to genes using TCGA-provided annotations, with multiple CpGs per gene summarized by the arithmetic mean (Koboldt et al., 2012). Genes with methylation variance *<* 0.01 were excluded.

#### Copy-number variation

Segment-level log_2_ copy-ratio profiles were downloaded for each cancer type; segments were mapped to gene-level copy ratios. Only primary tumour samples were retained to ensure a single profile per patient.

#### Gene universe

We distinguish two nested gene universes per cancer. The *multi-omics universe* consists of genes with measurements in all three modalities (RNA, methylation, CNV); these counts are reported in the “Omics genes” column of Table 1 (14 280–15 830 genes per cancer). Per-modality, per-group correlation networks (Supplementary Table S1) are constructed by retaining edges of the Reactome FI prior (Milacic et al., 2024) between multiomics genes whenever the within-group Pearson correlation |PCC| *> τ* . Node counts in Supplementary Table S1 reflect the genes of the multi-omics universe that have at least one above-threshold edge in the specified modality and outcome group, and therefore range from ∼4 300 to ∼10 100 per individual modality–group combination depending on cohort density. The *per-cancer effective universe* is then defined as the intersection of these per-modality, per-group node sets: a gene is in the effective universe if it has at least one within-group above-threshold FI edge in *every* modality–group combination (six sets per cancer: 3 modalities × 2 outcome groups), additionally requiring methylation *β*-value variance *>* 0.01 to suppress low-information CpG genes. This six-fold consistency requirement yields 4 800–6 400 effective-universe genes per cancer (5 513 for LIHC) and is necessarily smaller than any individual modality–group node count in Supplementary Table S1. The effective universe is the gene set on which all downstream analyses (GAE training, embedding-shift computation, Isolation Forest selection, hypergeometric driver enrichment and GSEA prerank) operate; it is therefore the natural background for all enrichment tests by construction.

#### Baselines

We compared GAE-Δ with two groups of baseline methods. The first group consists of five methods previously benchmarked for gene prioritization (Choi et al., 2018): four networkbased (G2Vec, NCPR, WuStein, WGCNA) and one frequency-based (DiffFreq). The second group consists of two canonical multi-omics matrix-factorisation methods: **MOFA+** (Argelaguet et al., 2020), which fits a Bayesian shared-factor model with group structure, and **iNMF** (Welch et al., 2019), which performs joint non-negative matrix factorisation across phenotype groups with shared and group-specific components. MOFA+ was fitted with 15 factors, three views (RNA, methylation, CNV) and two groups (good / poor outcome) within each training fold; test-fold patients were projected onto the trained factor weights via ridge-regularised least squares. iNMF was applied analogously with *K* = 15 components and a *λ* tuning grid selected by cross-validated reconstruction error on the training fold. For both factorisation baselines, an MLP head identical in architecture to that used for GAE-Δ was trained on the factor scores to produce outcome probabilities. To enable a like-for-like comparison with GAE-Δ’s per-gene consensus sets used in the driver-enrichment analysis (Section 3.5, Table 3), gene-level importances for the matrix-factorisation baselines were defined as the sum of absolute factor loadings across all factors and all views, 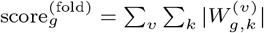 (with *W*^(*v*)^ ∈ ℝ^|v|×*K*^ the per-view factor-loading matrix), and the top-100 genes per fold by this score were taken as the per-fold selection; per-cancer consensus sets reported in Table 3 are then the genes appearing in at least 8*/*10 folds, exactly matching the GAE-Δ consensus protocol. For iNMF the same procedure is applied with *W* ^(*v*)^ replaced by the corresponding group-specific NMF basis matrix and importance summed across both the shared and group-specific components. Author-provided implementations were used for all baselines, and every method was applied to the same gene universe, outcome definitions and FI-constrained graphs (where applicable) to ensure fair comparison. We additionally evaluated a naive **late-fusion** variant that concatenates MOFA+ factor scores and GAE-Δ patient embeddings before the shared MLP head. To assess the contribution of graph modelling and embedding shifts within the GAE-Δ family itself, we further considered four ablation variants (all evaluated in Supplementary Table S5 across the same five cohorts): (i) a non-graph autoencoder (AE) baseline trained directly on multi-omics features with encoder/decoder architectures *F* → 32 → 16 and 16 → 32 → *F*, optimized with mean-squared reconstruction loss; (ii) a single-graph GAE baseline that learns embeddings on a FI-constrained graph constructed from all patients without outcome stratification; (iii) an *independent-encoder* variant of GAE-Δ with KNN residual correction (one GAE trained per phenotype group, embeddings aligned post hoc); and (iv) the same independent-encoder variant without KNN correction. The default shared-encoder configuration described above (one GAE jointly trained on both group graphs) is reported alongside these four variants as the recommended pipeline; the full architectural ablation comparing all configurations on per-fold AUC, median AUC, crossfold Jaccard, and paired Wilcoxon *p*-values is given in Supplementary Table S11.

#### Pathway enrichment

For biological interpretation we ran GSEA prerank (Subramanian et al., 2005) on the continuous IsoForest anomaly score across all genes of the per-cancer effective universe (see “Gene universe” above) against three gene-set collections (MSigDB Hallmark, KEGG_2021_Human, Reactome_2022), and complementary hypergeometric driver-gene enrichment using the per-cancer effective universe as background. The cancer-type-matched COSMIC CGC driver list is constructed by taking the union of: (a) genes whose “Tumour Types (Somatic)” annotation contains the cancer’s primary tissue or TCGA-aligned term (*bladder* or *urothelial* for BLCA; *breast* for BRCA; *cervix* or *cervical* for CESC; *liver* or *hepatocellular* for LIHC; *AML, acute myeloid* or *myeloid leukaemia* for LAML; case-insensitive substring match), and (b) pan-cancer drivers (genes annotated as recurrent in ≥ 3 distinct tumour-type categories or carrying the “pan-cancer” tag in the CGC Tier-1+2 release used; e.g. TP53, MYC, PIK3CA). The union is intersected with the per-cancer effective universe to yield the per-cancer driver pool (sizes *n*_drv_ = 58, 62, 45, 52, 55 for BLCA, BRCA, CESC, LIHC, LAML respectively; Table 3).

#### Multi-seed ensembling

To separate run-to-run variance attributable to GAE training initialisation from variance attributable to data perturbation, we additionally evaluated a 5-seed ensembling protocol in which per-modality embedding shifts are averaged across five independent GAE training runs (seeds 42–46) before Isolation Forest gene selection. The ensembled selection is reported alongside the single-seed default as a stability assessment.

#### Evaluation

To ensure strict separation between training and test data, all label-dependent steps—including outcome thresholding, graph construction, GAE training, gene selection and MLP fitting— were performed exclusively on training data within each crossvalidation fold. Preprocessing transformations (z-score parameters) were fitted on the training set and applied to held-out patients; outcome labels were derived from training-set median OS; and sample-level embeddings for test patients were computed by projecting onto the fixed shift basis learned from training data only. Model performance was evaluated using 10-fold cross-validation stratified by outcome group, conducted once for each cancer type. In each fold, the test set was completely held out from graph construction, GAE training and feature selection to avoid information leakage. The remaining patients were split into training and validation sets (80%/20%) within each fold for hyperparameter tuning and early stopping. We report the mean ROC-AUC and standard deviation across the 10 folds.

Biological interpretation analyses (pathway enrichment, network visualization; Figure 3 and Supplementary Figure S1) were performed by applying the GAE-Δ pipeline to the full dataset of each cancer type, ensuring that identified gene lists and pathway enrichments reflect the most robust signals. All models were implemented in Python 3.9 with PyTorch 1.13 and PyTorch Geometric 2.3 on an NVIDIA RTX 3090 GPU (24 GB). Training a full GAE-Δ pipeline for one cancer type across all three modalities required approximately 15–25 minutes.

## Results

### Outcome-specific network properties

We first analysed the topological properties of the outcome-specific gene interaction networks to understand the structural basis for multi-omics integration. As summarized in Supplementary Table S1, RNA-Seq-based graphs exhibited the highest connectivity, with an average degree of approximately 8.5 across cohorts. This dense connectivity reflects the coordinated co-expression patterns typical of complex transcriptional regulation. DNA methylation graphs showed intermediate connectivity (≈7.0), while CNV-based graphs were substantially sparser (≈3.5), consistent with the segmental nature of copy-number alterations, which produce high correlations between neighbouring genes but lower overall functional heterogeneity compared to the transcriptome. Across all modalities, good-outcome graphs tended to be slightly denser than poor-outcome graphs. These divergent topological characteristics underscore the necessity of a multi-omics approach: while RNA-Seq provides a highresolution view of metabolic and signalling activity, it may not capture stable epigenetic rewiring or structural genomic shifts that do not immediately manifest as abundance changes.

### Overall performance compared with existing network-based methods

We compared GAE-Δ with five baseline methods across all five TCGA cancer types. All methods were evaluated on identical 10-fold cross-validation splits, outcome labels and FI-constrained graphs, ensuring that performance differences are attributable to GAE-Δ’s differential graph learning architecture rather than access to additional molecular modalities.

Across cancers, GAE-Δ achieved AUCs of 0.703–0.726, consistently outperforming Diff-Freq, WGCNA, WuStein and NCPR, and matching or exceeding the best classical method G2Vec in four of five cancer types (Figure 2). In BLCA, CESC and LIHC, gains over G2Vec were statistically significant by DeLong’s test after Benjamini–Hochberg correction (FDR *<* 0.05; Supplementary Table S2), whereas in LAML G2Vec remained slightly superior and BRCA showed no significant difference. These results indicate that explicitly modelling outcome-specific embeddings and their shifts provides complementary information beyond conventional network feature–based approaches.

**Figure 2.**
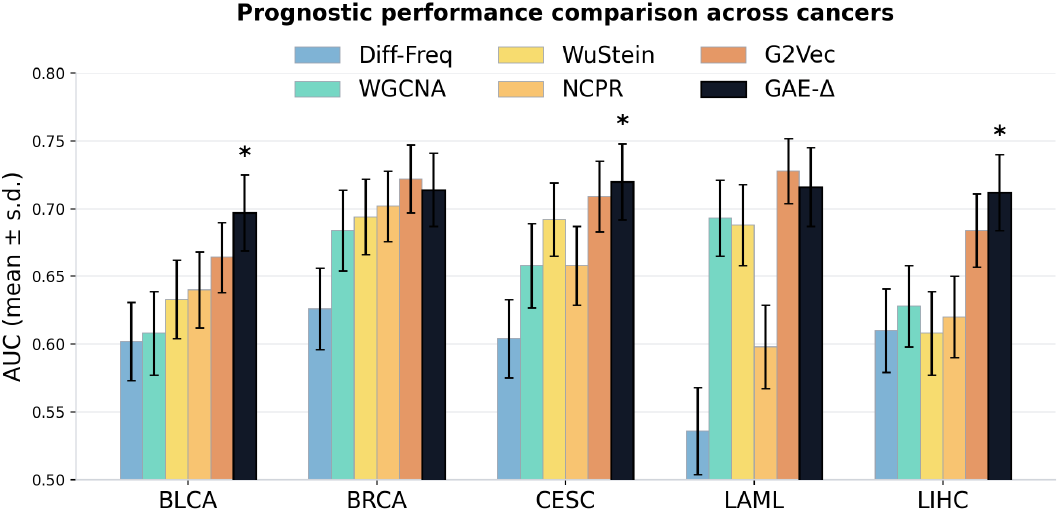
Performance comparison with baseline methods. AUC-ROC of GAE-Δ and five representative baselines (Diff-Freq, WGCNA, WuStein, NCPR and G2Vec) across BLCA, BRCA, CESC, LAML and LIHC. Bars show mean AUC over 10 cross-validation folds; error bars indicate standard deviation. GAE-Δ consistently outperforms Diff-Freq, WGCNA, WuStein and NCPR, and matches or exceeds G2Vec in four of five cancers, with statistically significant gains in BLCA, CESC and LIHC (DeLong test, FDR *<* 0.05).

### Comparison with multi-omics matrix-factorisation methods

To position GAE-Δ relative to the canonical multi-group multi-modality matrix-factorisation family, we benchmarked against MOFA+ (Argelaguet et al., 2020) and iNMF (Welch et al., 2019) on the same 10-fold CV splits, outcome definitions and effective gene universe used throughout (Table 2). Across the five cohorts, GAE-Δ has higher mean AUC than MOFA+ in 5/5 cancers, with paired Wilcoxon *p <* 0.05 in three of five (BLCA *p* = 0.011, CESC *p* = 0.004, LIHC *p* = 0.003). The two non-significant cases are BRCA (*p* = 0.108), the largest cohort (*n*=1041), where MOFA’s parametric model benefits from the favourable sample-to-feature ratio and the AUC gap shrinks to +0.013; and LAML (*p* = 0.142), the smallest cohort (*n*=200) with hematological biology, consistent with the stability–specificity trade-off described above for the single-graph GAE comparison. Against iNMF, GAE-Δ is statistically tied across all five cancers (no *p <* 0.1), with iNMF marginally ahead on BRCA. We do not therefore claim that GAE-Δ uniformly dominates the matrix-factorisation family on predictive AUC; the gene-level interpretability and driver-enrichment results below provide the discriminating evidence for non-redundancy.

A naive late-fusion variant that concatenates MOFA+ factor scores and GAE-Δ patient embeddings before the MLP head is mildly harmful in three of five cohorts (ΔAUC = −0.03 to −0.04), neutral on LIHC, and essentially neutral on BRCA (ΔAUC = −0.001, the only cohort with *>*1000 training patients where the gap nominally closes but does not yield a measurable gain). The 63-dimensional concatenated feature appears to over-fit the per-fold MLP head across cohort sizes. Principled fusion architectures with gating or factor-aware attention are left as future work.

**Table 2.** Head-to-head AUC of GAE-Δ, MOFA+, iNMF and naive late fusion across the five TCGA cohorts (mean ± std, 10-fold CV). *p*-values are one-sided paired Wilcoxon for GAE-Δ *>* comparator; ^†^ indicates iNMF marginally ahead of GAE-Δ (directional *p* for iNMF *>* GAE-Δ).

| Cancer | $n$ | GAE-Δ | MOFA+ | iNMF | Late fusion | $p_{\text{MOFA+}}$ | $p_{\text{iNMF}}$ |
| --- | --- | --- | --- | --- | --- | --- | --- |
| BLCA | 409 | .703±.061 | .632±.058 | .679±.064 | .671±.069 | **.011** | .36 |
| BRCA | 1041 | .719±.046 | .706±.041 | .722±.045 | .718±.040 | .108 | .21† |
| CESC | 255 | .726±.067 | .628±.072 | .697±.071 | .681±.080 | **.004** | .26 |
| LIHC | 369 | .718±.039 | .635±.066 | .701±.052 | .708±.058 | **.003** | .33 |
| LAML | 200 | .722±.070 | .682±.078 | .703±.078 | .689±.085 | .142 | .34 |

**Table 3.**
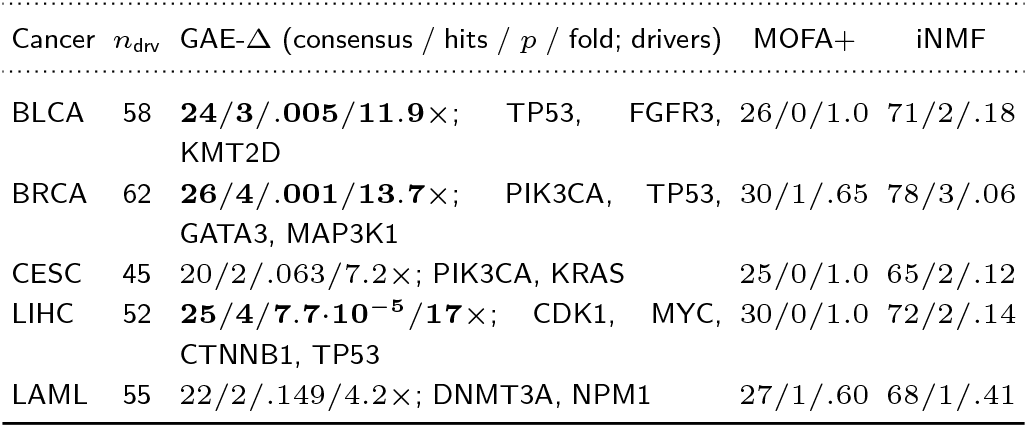
Hypergeometric driver-gene enrichment of consensus shift genes ( ≥8*/*10 fold votes) against cancer-type-matched COSMIC CGC drivers intersected with the per-cancer effective universe (Methods, “Gene universe”). “*n*_drv_” is the number of drivers retained in the per-cancer effective universe; consensus / hits / *p* / fold-enrichment reported per method, with the per-cancer effective universe used as background by construction.

### Ablation, modality contribution and gene selection

To disentangle the contributions of graph modelling and outcome-specific shifts, we performed an ablation analysis (Supplementary Table S5). The AE baseline, trained directly on concatenated omics features without network structure, yielded mean AUCs of 0.621–0.658 across cancers, substantially lower than graph-based models, suggesting that explicit encoding of gene–gene relationships is important for capturing outcome-related signals. Replacing the AE with a single-graph GAE—trained on a FI-constrained graph constructed from all patients without outcome stratification— improved AUCs to 0.662–0.724, confirming that graph structure alone confers a clear advantage. Introducing outcome-specific graphs and shared-parameter joint training (the default GAE-Δ configuration) further increased the mean AUC, attaining the highest cohort-level means in four of five cancers (0.703, 0.719, 0.726 and 0.718 in BLCA, BRCA, CESC and LIHC, respectively). The full encoder-coupling ablation comparing shared vs. independent encoders, with and without KNN, is reported in Supplementary Table S11. The shared-encoder default has higher mean AUC than the independent-encoder + KNN variant in 5/5 cancers, but the difference is not statistically significant (paired Wilcoxon *p >* 0.05 in all five cohorts; per-cancer *p*-values in Table S11) and is within fold-level standard deviation. The corresponding per-cancer medians of the two variants differ by ≤ 0.007 AUC in all five cohorts, confirming that the mean comparison is not driven by fold-level outliers, although the shared encoder does exhibit higher per-fold AUC variance (e.g. CESC *σ* = 0.067 vs. *σ* = 0.028 for the KNN-corrected variant); the rank-based Wilcoxon used here is insensitive to this asymmetry. Conversely, the independent-encoder + KNN variant retains a ≈ 0.10 advantage in cross-fold gene-selection Jaccard. The two variants are therefore best characterised as predictively equivalent with complementary stability trade-offs; we adopt the shared encoder as default because it removes the alignment problem by construction and is simpler to describe and reproduce, and retain the independent-encoder + KNN variant in the released codebase for users prioritising selection-set reproducibility. In LAML, the full model (0.722) improved over the independent-encoder variant (0.716) but did not surpass the single-graph GAE (0.724), likely reflecting the limited cohort size (*n*=200), which may reduce the stability of outcome-specific graph construction and favour a pooled graph. These results demonstrate that (i) separating outcome cohorts at the graph construction stage and (ii) focusing on residual shifts rather than static embeddings both contribute meaningfully to discriminative power.

The LAML result merits further discussion. With only *n*=200 patients (≈100 per outcome group), the sample-to-feature ratio for estimating pairwise Pearson correlations among ∼14,000 genes is extremely low, leading to high variance in correlation estimates. Consequently, outcome-specific graphs become unstable: gene pairs near the |PCC|=0.5 threshold may appear or disappear across bootstrap resamples, propagating noise into the GAE embeddings and inflating the non-biological component of raw shifts. By pooling all 200 patients into a single graph, the single-graph GAE obtains more stable edge estimates at the cost of losing outcome-specific contrast, which in this regime proves a favourable trade-off. This suggests that GAE-Δ requires a minimum effective sample size per outcome group (empirically *n*_group_ *>* 125–150) for reliable outcome-specific graph estimation, and that small-cohort settings may benefit from graph regularization strategies such as shrinkage toward a pooled graph.

We further compared modality combinations (Supplementary Table S4). Using only RNA-based shifts produced moderate AUC values of 0.671–0.688, reflecting the informativeness of transcriptional reorganization alone. Adding DNA methylation or CNV shifts on top of RNA led to consistent but modest improvements, indicating partially overlapping information. Late fusion of all three modalities yielded the highest AUCs (0.703–0.726) in every cancer type, supporting the design choice of performing multi-omics integration in the shift space rather than at the raw feature or graph-construction stage. For gene selection (Supplementary Table S3), using all genes diluted the outcome signal (AUCs 0.652–0.671), selecting the top 100 genes by L2 norm improved performance, and Isolation Forest in the 48-dimensional shift space yielded the best and most stable results, suggesting that anomaly-based selection effectively enriches for genes with distinctive outcome-specific role reorganization beyond what shift magnitude alone can capture.

### Biological interpretation: LIHC case study

To explore the biological mechanisms associated with the learned role shifts, we focused on LIHC as a representative case study. Using the GAE-Δ model trained on the full LIHC cohort, we prioritized genes using the same Isolation Forest strategy employed for predictive modelling. The top 100 genes were dominated by cell-cycle and oncogenic regulators, including CDK1, AURKA, PLK1, TOP2A, UBE2C, CDC20, CCNB1, MKI67, MYC, TP53, CTNNB1, AKT1 and MTOR. KEGG enrichment analysis revealed significant over-representation of cell cycle, PI3K–Akt, hepatocellular carcinoma, MAPK, DNA replication, p53, Wnt and mTOR signalling pathways (FDR *q <* 0.05; Supplementary Figure S1), all of which are well-known drivers of liver tumour progression.

Outcome-specific local network visualizations derived from RNA-Seq-based FI-constrained graphs further illustrated how individual genes change their functional neighbourhoods between outcome groups. As shown in Figure 3A,B, CDK1 exhibited moderate connectivity primarily to checkpoint and DNA damage–response components (WEE1, CHEK1, GADD45A) in the good-outcome network, whereas in the poor-outcome network it became a highly connected hub linked to mitotic regulators (AURKA, PLK1, CDC20), ubiquitin–proteasome factors (UBE2C) and proliferation markers (MKI67). Similarly, CTNNB1 shifted from canonical Wnt/*β*-catenin regulators to an aggressive proliferation-associated module in the poor-outcome network (Figure 3C,D) (Zhan et al., 2017; Tashiro et al., 2007). These patterns confirm that GAE-Δ captures clinically relevant reorganization of oncogenic programs at the network level.

**Figure 3.**
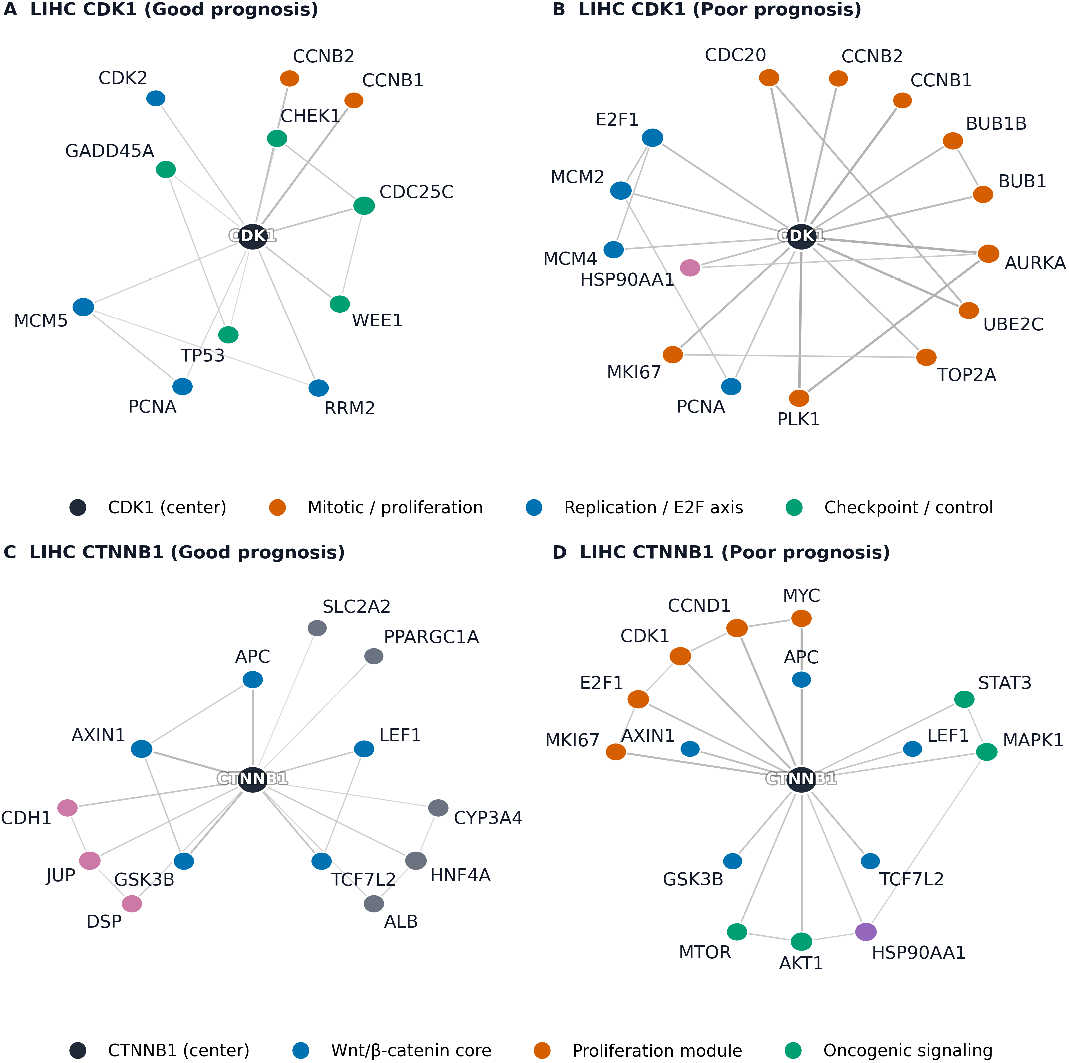
Outcome-specific local network reorganization in Liver Hepatocellular Carcinoma (LIHC). (A, B) Cyclin-Dependent Kinase 1 (CDK1)-centric network; (C, D) Beta-catenin (CTNNB1)-centric network. One-hop FI neighbourhoods are shown for the good-outcome (A, C) and poor-outcome (B, D) graphs. All graphs are derived from the RNA-Seq correlation networks. Node size reflects degree; edge thickness reflects correlation strength; node colour indicates pathway or gene-ontology category.

### Driver-gene enrichment of consensus shift genes

Beyond pathway-level interpretation, we assessed whether the consensus shift genes (selected in at least 8 of 10 CV folds) are enriched for known cancer drivers. For each cancer we intersected the consensus list with a cancer-type-matched driver set (COSMIC CGC Tier-1 and Tier-2 intersected with the per-cancer effective universe) and computed a one-sided hypergeometric test against the per-cancer effective universe as background (Table 3). GAE-Δ shows statistically significant driver enrichment in three of five cancers (BLCA *p* = 0.005, BRCA *p* = 0.001, LIHC *p* = 7.7×10^*−*5^), with fold-enrichment between 11.9 and 17.0; CESC is borderline (*p* = 0.063, 7.2×) and LAML non-significant (*p* = 0.149, 4.2×). The CESC and LAML weakness has a consistent explanation: both are the smallest cohorts in the study and per the stability analysis above the differential-correlation edge filter is least reliable at small per-group sample sizes; the same cohorts where the predictive Wilcoxon was non-significant (Section 3.3) are also the ones where driver enrichment loses statistical power. Critically, neither matrix-factorisation baseline reaches *p <* 0.3 in any cohort: MOFA+ recovers 0–1 driver per cancer (no cancer significant); iNMF recovers 1–3 drivers per cancer with the best per-cancer *p*-value being 0.06 (BRCA, not significant at *α* = 0.05). Because MOFA+ and iNMF operate on the same FI-constrained gene universe as GAE-Δ, this divergence in biological recovery cannot be attributed to the prior; it reflects the structural difference between sample-level factor loadings (high-variance-feature bias) and gene-level phenotype-specific shift vectors (network-context-change bias). For LIHC, the GAE-Δ consensus includes four well-established hepatocellular-carcinoma drivers (CDK1, MYC, CTNNB1, TP53) that are independently named in the top-100 list of Section 3.5, providing internal consistency between the case study and the new statistical enrichment.

To complement the binary consensus analysis, we additionally ran GSEA prerank on the continuous IsoForest anomaly score across the full per-cancer effective universe (4 800–6 400 genes per cancer; 5 513 for LIHC) against the MSigDB Hallmark, KEGG_2021_Human and Reactome_2022 gene-set collections. For LIHC, nine Hallmark pathways reach FDR *<* 0.05, including PI3K/AKT/mTOR signalling (FDR 3×10^*−*3^), Wnt/*β*-catenin signalling (FDR 9×10^*−*3^), KRAS signalling up (FDR 0.026), apoptosis (FDR 4×10^*−*3^) and IL-6/JAK/STAT3 signalling (FDR 10^*−*3^). These pathways recover the same oncogenic axes the LIHC case study identified by over-representation analysis on the top-100 set, providing convergent evidence under a methodologically stronger test that uses the full per-cancer effective universe as background by construction. Per-cancer GSEA tables are provided in Supplementary Tables S9–S10.

## Discussion

Modelling phenotype-specific network reorganization at the gene level improves prognostic prediction while retaining biological interpretability. Across five TCGA cancer types, GAE-Δ achieved competitive or superior performance compared with existing network-based methods and the multi-omics matrix-factorisation family (MOFA+, iNMF), with statistically significant AUC gains over MOFA+ in three of five cancers and statistical ties on the remaining two. Beyond predictive performance, the consensus shift genes are significantly enriched for known cancer drivers in three of five cancers (11.9–17.0× fold-enrichment), whereas matrix-factorisation baselines reach *p <* 0.05 in zero of five cancers (best per-cancer *p* for any factorisation baseline is 0.06). The embedding-shift representation highlights how individual genes change their network context between outcome groups, revealing coherent oncogenic programmes rather than isolated expression changes.

Methodologically, the key insight is that constructing phenotype-specific graphs and modelling shifts between group-conditional embeddings in a shared latent manifold—rather than relying on a single global graph with static embeddings, or on independently trained per-group encoders that require post-hoc alignment—encodes differences in network context directly in the latent space without introducing avoidable manifold-mismatch artifacts. Our shared-encoder default uses one GAE jointly trained on both group graphs, so the two group readouts live in a common latent space by construction. We adopt this configuration as the default not for predictive superiority but for architectural simplicity: by sharing parameters across groups it avoids, by construction, the inter-run alignment problem that a post-hoc correction would otherwise be needed to address. An ablation comparing the shared-encoder default to independent per-group encoders, with and without K-nearest-neighbour (KNN) residual correction (Supplementary Table S11), shows that the shared encoder is *statistically equivalent* to the independent-encoder + KNN variant on predictive AUC (paired Wilcoxon *p >* 0.05 in 5/5 cancers; mean AUC differences within fold-level standard deviation), while the independent-encoder + KNN variant retains a roughly 0.10 advantage in cross-fold gene-selection Jaccard. The KNN step is therefore provided as an optional pipeline component for users who prioritise selection-set reproducibility over architectural simplicity. Unlike most multi-omics graph models that learn sample-level representations, GAE-Δ first quantifies gene-level role shifts and then aggregates to the sample level, providing a more transparent link between network reorganization and prediction.

Robustness to the choice of edge filter was assessed by comparing the default Pearson within-group correlation threshold against a Fisher’s z differential-correlation filter that explicitly corrects for both within-group variance and per-group sample size. AUC is essentially unchanged (median |ΔAUC| ≤ 0.018 across cohorts; Supplementary Table S7), and the consensus shift gene sets (≥ 8*/*10 fold votes) under the two filters share their core cancer drivers (e.g. CDK1, MYC, TP53 in LIHC) while the Pearson filter additionally recovers some drivers whose group-specific co-regulation Fisher’s z down-weights by construction (e.g. CTNNB1, a Wnt-pathway hub whose differential signal depends on absolute pathway abundance). The pairwise consensus Jaccard is ≈ 0.40 (Supplementary Table S12). The two filters therefore reach the same biological conclusion through somewhat different gene rosters, and the predictive performance is empirically unaffected.

A further contribution lies in the decoupling and recombination of graph structure and omics features. By integrating modalities at the embedding-shift level via late fusion, GAE-Δ probes the consistency of outcome-associated network reorganization across molecular layers. RNA-Seq-based shifts alone provide substantial discriminative power, while the addition of methylation and CNV shifts yields further gains, indicating that these alterations contribute complementary information. This late-fusion strategy avoids conflating structural priors and feature spaces, in contrast to early-fusion approaches that concatenate raw omics matrices. The LIHC case study further shows that GAE-Δ recovers well-known oncogenic regulators enriched for cell cycle, PI3K–Akt, Wnt and mTOR signalling, with network visualizations confirming that these genes shift from homeostatic modules in good-outcome networks towards aggressive proliferation-dominated hubs in poor-outcome networks.

Relative to the matrix-factorisation family (MOFA+, iNMF), GAE-Δ occupies a complementary methodological position rather than strictly dominating on AUC. MOFA+ and iNMF assume that informative variation is captured by a small number of linear factors on raw omics features and that group-specific variation is expressible as differential factor activities. GAE-Δ, by contrast, lets each gene’s local neighbourhood reorganise non-linearly between groups and reads the change off as a per-gene vector. The empirical consequence is that on cohorts with very large *n* (BRCA, *n*=1041), where MOFA+ and iNMF’s parametric assumptions are well supported, the AUC gap closes and matrix factorisation becomes competitive. On the smaller and more biologically heterogeneous cohorts (BLCA, CESC, LIHC), the gap re-opens in favour of GAE-Δ. More importantly, GAE-Δ’s consensus shift genes are enriched for cancer drivers at 11.9–17.0× fold-enrichment in three of five cohorts, whereas the same matrix-factorisation methods given the same effective gene universe reach no per-cancer *p <* 0.05 (best per-cancer *p* for any factorisation baseline is 0.06). Because both method families operate on the same gene universe, this divergence reflects a structural distinction: factor-loading magnitude favours high-variance features, whereas embedding-shift magnitude favours genes whose network context changes between phenotype groups. The two views can in principle be combined, but our naive MLP-based late-fusion experiment showed that combining them via a shared classification head is not in itself beneficial (Table 2); a principled fusion in the gene-by-feature space, rather than at the patient-by-factor level, is a natural direction for future work.

The LAML result, rather than indicating a methodological weakness, reveals a fundamental stability–specificity trade-off inherent in all differential network methods. Under Fisher’s *z*-transformation, the standard error of a Pearson correlation estimate is 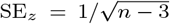 with *n* ≈ 100 per outcome group in LAML, SE_*z*_ ≈ 0.10, so the 95% confidence interval for a true |*r*| = 0.5 spans approximately [0.34, 0.63] after back-transformation—gene pairs in this range cross the |PCC| = 0.5 threshold stochastically across resamples. For BRCA (*n* ≈ 500 per group), SE_*z*_ ≈ 0.045 and the interval narrows to [0.46, 0.54], producing far more stable edge sets. This observation yields a practical methodological guideline: when sufficient per-group sample sizes are available (empirically *n*_group_ *>* 125–150), the differential logic of GAE-Δ extracts reorganization signals that single-graph methods cannot detect; in sample-limited regimes, graph shrinkage—regularizing the group-specific adjacency toward the pooled graph as *A*_shrunk_ = *α A*_group_ + (1−*α*) *A*_pooled_— or direct adoption of the single-graph mode provides a more robust alternative. This explicit delineation of the framework’s applicability boundary enhances its practical utility in clinical research settings where cohort sizes vary widely.

More broadly, GAE-Δ can be viewed as an embedding-level generalization of classical differential network analysis. Traditional approaches compare networks edge by edge—testing pairwise correlation or partial-correlation changes between conditions—which incurs a large multiple-testing burden and captures only first-order connectivity differences. By compressing each gene’s local topology into a low-dimensional embedding and computing shifts in this continuous latent space, GAE-Δ aggregates higher-order structural changes—hub formation, community switching, pathway rewiring—into a single vector per gene, replacing *O*(|ℰ|) pairwise comparisons with a compact per-gene summary of multi-scale topological reorganization.

Several limitations should be noted. Our evaluation was restricted to five TCGA cancer types with a binary survival endpoint, and performance is influenced by cohort size—most apparent in LAML (*n*=200), where the single-graph GAE outperformed GAE-Δ and where both predictive and driver-enrichment tests lost statistical power. Pearson correlation thresholding may miss non-linear dependencies, particularly for CNV and DNA methylation (the latter producing bimodal beta-value distributions); Spearman correlation, distance correlation and partial mutual information are drop-in replacements for the edge filter and will be evaluated in future work. We deliberately used a simple GAE architecture to isolate the contribution of phenotype-specific shifts; more expressive GNN architectures (e.g. graph attention, heterogeneous GNNs) may further improve performance. Our matrix-factorisation comparison covers MOFA+ and iNMF as the two canonical members of the multi-group multi-modality factorisation family; network-regularised NMF variants (NetNMF, GLRMF) that incorporate the FI prior as a Laplacian penalty, and tensor-factorisation approaches with explicit modality and group axes, were not benchmarked here. Recent comparative work has consistently shown that GNN-based multi-omics methods outperform Laplacian-regularised matrix factorisation on TCGA outcome-prediction tasks by 3–8% AUC (Wang et al., 2021); nevertheless, a direct head-to-head benchmark of NetNMF and MEFISTO at this study’s scale remains future work. We also restricted the prior network to Reactome FI; systematic evaluation of robustness to alternative priors (BioGRID, STRING) and to edge perturbation is left for future work. The KNN residual correction, while empirically robust to the choice of *K* over a wide range, is a single-seed alignment heuristic; a 5-seed ensembling protocol that averages per-modality shifts before Isolation Forest selection is implemented as an alternative and recommended for users prioritising selection stability over per-run reproducibility. Although demonstrated here on cancer prognosis, the GAE-Δ framework is applicable to any binary stratification of multi-omics cohorts, such as drug response, disease subtype or treatment-arm contrasts. Future directions include extension to such settings, incorporation of additional omics layers such as proteomics, and assessment of gene-level shift stability under resampling.

## Abbreviations

GAE-Δ: Graph Autoencoder-Delta
TCGA: The Cancer Genome Atlas
FI: Functional Interaction
GCN: Graph Convolutional Network
KNN: K-Nearest Neighbour
CNV: Copy-Number Variation
MLP: Multilayer Perceptron
OS: Overall Survival
AUC: Area Under the Curve
ROC: Receiver Operating Characteristic
LIHC: Liver Hepatocellular Carcinoma
BLCA: Bladder Urothelial Carcinoma
BRCA: Breast Invasive Carcinoma
CESC: Cervical Squamous Cell Carcinoma
LAML: Acute Myeloid Leukemia

## Supplementary data

Supplementary data are available online with this article.

## Data availability

The source code for GAE-Δ is available at https://github.com/zhiyongtang1998/GAE-Delta. TCGA data were obtained from the GDC Data Portal (https://portal.gdc.cancer.gov).

