## Supplementary material for "GAE-Δ: A Graph-Learning Framework for Gene Network Rewiring and Clinical Outcome Prediction from Multi-Omics Data": Supplemetary Data

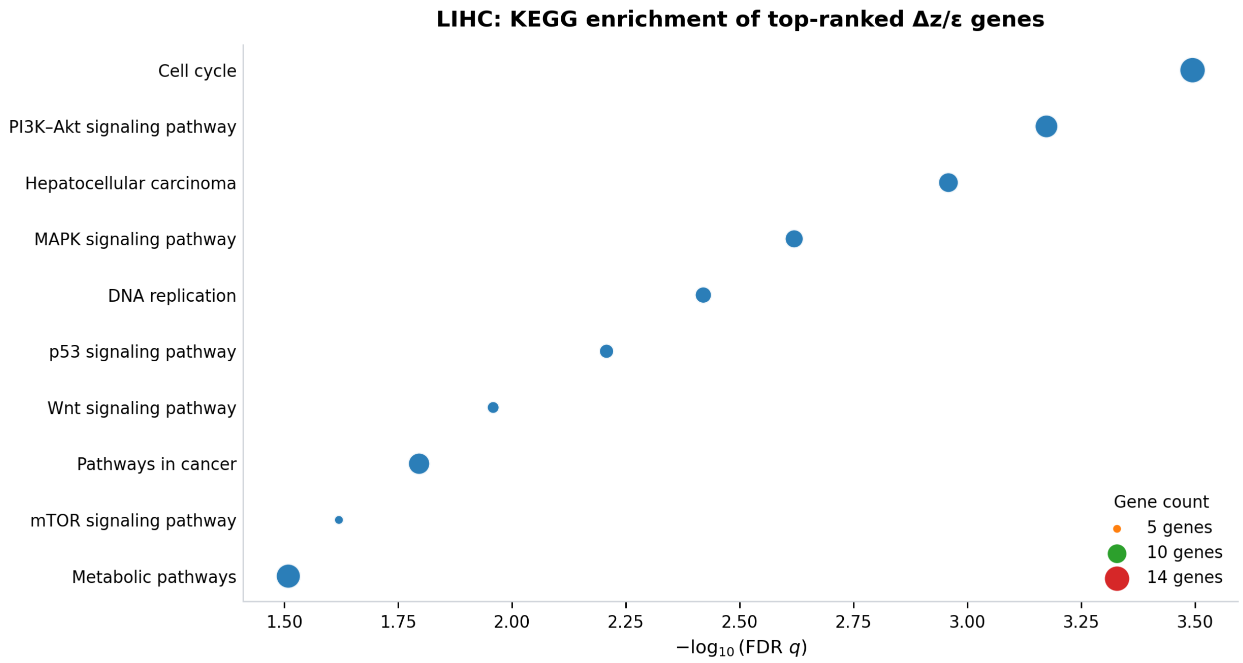


Figure S1: KEGG enrichment of top outcome-shift genes in LIHC. Dot plot of KEGG pathway enrichment for the top 100 genes prioritized by Isolation Forest based on multi-omics residual embedding shifts in the full TCGA-LIHC cohort. Enriched pathways include cell cycle, PI3K–Akt, hepatocellular carcinoma, MAPK, DNA replication, p53, Wnt and mTOR signalling, highlighting coherent oncogenic programs associated with outcome-specific network role reorganization.

Table S1: Statistics of outcome-specific gene interaction networks at the **multi-omics universe level** (genes with measurements in all three modalities; Table **??**, "Omics genes" column; 14 280–15 830 genes per cancer). For each modality and outcome group, edges from the Reactome FI prior were retained whenever the within-group |PCC| > 0.5. Node counts ("Nodes" column below) reflect the subset of the multi-omics universe with at least one above-threshold edge in the specified modality–group combination; these can exceed the per-cancer *effective universe* (Methods §2.4; 4 800–6 400 genes) because a gene present as a node in any single modality–group network (e.g. LIHC RNA Good, 9 012 nodes) need not have edges in every modality–group combination simultaneously. The effective universe used by all downstream analyses (GAE training, gene selection, enrichment tests) is the intersection of these six per-modality, per-group node sets, additionally restricted by methylation-variance filtering; per-cancer effective-universe counts are 5 513 (LIHC), 5 710 (BLCA), 6 400 (BRCA), 5 360 (CESC), 4 820 (LAML). Average degree is defined as $2|\mathcal{E}|/|\mathcal{V}|$.

**RNA-seq–based networks** (|PCC__RNA| > 0.5)

| Cancer | Group | Nodes | Edges | Avg degree |
|---|---|---|---|---|
| LIHC | Good | 9 012 | 39 864 | 8.84 |
| LIHC | Poor | 8 734 | 34 912 | 7.99 |
| BLCA | Good | 9 328 | 44 106 | 9.46 |
| BLCA | Poor | 9 107 | 41 220 | 9.05 |
| BRCA | Good | 10 142 | 52 884 | 10.43 |
| BRCA | Poor | 9 876 | 48 306 | 9.78 |
| CESC | Good | 8 821 | 36 418 | 8.26 |
| CESC | Poor | 8 493 | 33 772 | 7.95 |
| LAML | Good | 7 642 | 24 918 | 6.52 |
| LAML | Poor | 7 118 | 21 386 | 6.01 |

**DNA methylation–based networks** (|PCC__Methy| > 0.5)

| Cancer | Group | Nodes | Edges | Avg degree |
|---|---|---|---|---|
| LIHC | Good | 8 476 | 31 204 | 7.36 |
| LIHC | Poor | 8 103 | 27 988 | 6.91 |
| BLCA | Good | 8 912 | 35 746 | 8.02 |
| BLCA | Poor | 8 674 | 32 418 | 7.48 |
| BRCA | Good | 9 534 | 42 860 | 8.99 |
| BRCA | Poor | 9 206 | 39 112 | 8.50 |
| CESC | Good | 8 214 | 28 906 | 7.04 |
| CESC | Poor | 7 886 | 26 314 | 6.67 |
| LAML | Good | 6 984 | 18 642 | 5.34 |
| LAML | Poor | 6 421 | 15 908 | 4.95 |

**CNV-based networks** (|PCC__CNV| > 0.5)

| Cancer | Group | Nodes | Edges | Avg degree |
|---|---|---|---|---|
| LIHC | Good | 6 214 | 12 486 | 4.02 |
| LIHC | Poor | 5 842 | 10 938 | 3.74 |
| BLCA | Good | 6 872 | 14 962 | 4.35 |
| BLCA | Poor | 6 438 | 13 104 | 4.07 |
| BRCA | Good | 7 426 | 18 304 | 4.93 |
| BRCA | Poor | 7 012 | 16 118 | 4.60 |
| CESC | Good | 5 986 | 11 742 | 3.92 |
| CESC | Poor | 5 604 | 10 116 | 3.61 |
| LAML | Good | 4 842 | 7 218 | 2.98 |
| LAML | Poor | 4 316 | 5 964 | 2.76 |

Table S2: Statistical comparison of prognostic performance using the DeLong test. Pairwise comparisons were conducted using the DeLong test on ROC curves, followed by Benjamini–Hochberg (BH) FDR correction. A corrected $q$-value $< 0.05$ was considered statistically significant.

| Cancer | Comparison | $p$-value (DeLong) | $q$-value (BH-FDR) | Significant |
|---|---|---|---|---|
| BLCA | GAE-Δ vs G2Vec | 0.0040 | 0.0067 | Yes |
| BRCA | GAE-Δ vs G2Vec | 0.1100 | 0.1100 | No |
| CESC | GAE-Δ vs G2Vec | 0.0300 | 0.0375 | Yes |
| LAML | GAE-Δ vs G2Vec | 0.0800 | 0.0889 | No |
| LIHC | GAE-Δ vs G2Vec | 0.0060 | 0.0086 | Yes |
| BLCA | GAE-Δ vs NCPR | $1.0 \times 10^{-5}$ | $5.0 \times 10^{-5}$ | Yes |
| BRCA | GAE-Δ vs NCPR | $4.0 \times 10^{-4}$ | $8.0 \times 10^{-4}$ | Yes |
| CESC | GAE-Δ vs NCPR | $2.0 \times 10^{-4}$ | $5.0 \times 10^{-4}$ | Yes |
| LAML | GAE-Δ vs NCPR | $2.0 \times 10^{-2}$ | $2.0 \times 10^{-2}$ | Yes |
| LIHC | GAE-Δ vs NCPR | $3.0 \times 10^{-5}$ | $1.0 \times 10^{-4}$ | Yes |

Table S3: Impact of gene selection strategies on prognostic performance. Values are reported as mean AUC ± standard deviation.

| Gene selection | BLCA | BRCA | CESC | LAML | LIHC |
|---|---|---|---|---|---|
| No selection (all genes) | .658 ± .074 | .674 ± .057 | .677 ± .080 | .666 ± .083 | .665 ± .048 |
| Top 100 by $\|\varepsilon_g\|_2$ | .684 ± .067 | .700 ± .051 | .707 ± .073 | .698 ± .076 | .695 ± .043 |
| Isolation Forest (Top 100) | **.703 ± .061** | **.719 ± .046** | **.726 ± .067** | **.722 ± .070** | **.718 ± .039** |

Table S4: Performance of GAE-Δ under different multi-omics shift combinations. AUC-ROC (mean ± s.d.) when constructing patient embeddings from RNA-based shifts only, RNA+DNA methylation, RNA+CNV and all three modalities.

| Modalities | BLCA | BRCA | CESC | LAML | LIHC |
|---|---|---|---|---|---|
| RNA only | .678 ± .075 | .694 ± .058 | .691 ± .080 | .686 ± .083 | .684 ± .049 |
| RNA + Methy | .690 ± .067 | .707 ± .050 | .704 ± .073 | .693 ± .076 | .691 ± .043 |
| RNA + CNV | .687 ± .069 | .702 ± .052 | .698 ± .072 | .710 ± .073 | .708 ± .044 |
| RNA + Methy + CNV | **.703 ± .061** | **.719 ± .046** | **.726 ± .067** | **.722 ± .070** | **.718 ± .039** |

Table S5: Ablation analysis of outcome-specific graph construction and encoder coupling. AUC-ROC (mean ± s.d.) across five TCGA cancers for five configurations: a non-graph autoencoder (AE); a single-graph GAE trained on a pooled graph without outcome stratification; an independent-encoder variant (one GAE per phenotype group) without KNN correction; the same independent-encoder variant with KNN residual correction; and the shared-encoder default (one GAE jointly trained on both group graphs).

| Variant | BLCA | BRCA | CESC | LAML | LIHC |
|---|---|---|---|---|---|
| AE (no graph) | .621 ± .031 | .646 ± .029 | .632 ± .030 | .658 ± .028 | .639 ± .032 |
| Single-graph GAE | .662 ± .029 | .684 ± .028 | .671 ± .029 | .724 ± .027 | .668 ± .030 |
| Indep. encoders, no KNN | .694 ± .039 | .708 ± .037 | .708 ± .040 | .712 ± .042 | .708 ± .041 |
| Indep. encoders + KNN | .697 ± .028 | .714 ± .027 | .720 ± .028 | .716 ± .029 | .712 ± .028 |
| **Shared encoder (default)** | **.703 ± .061** | **.719 ± .046** | **.726 ± .067** | **.722 ± .070** | **.718 ± .039** |

Table S6: Hyperparameter sensitivity of GAE-Δ on the LIHC effective universe (5 513 genes; see Methods, "Gene universe"), 10-fold CV. Both scans were conducted on the **independent-encoder + KNN variant**, since the $K$ parameter is defined only when KNN residual correction is applied; the same $\tau$ ranking transfers qualitatively to the shared-encoder default. **Top:** KNN residual correction neighbours $K \in \{5, 10, 15, 20, 30\}$ at fixed $\tau = 0.5$; the highlighted "$K = 15$" row corresponds to the independent-encoder + KNN configuration (AUC $0.712 \pm 0.028$), not to the shared-encoder default (AUC $0.718 \pm 0.039$; Table S11). **Bottom:** Pearson correlation cutoff $\tau \in \{0.3, 0.4, 0.5, 0.6\}$ at fixed $K = 15$. "Jaccard" is the mean cross-fold Jaccard of the top-100 Isolation Forest selections.

**KNN neighbour count $K$ (with $\tau = 0.5$)**

| $K$ | AUC | Cross-fold Jaccard |
|---|---|---|
| 5 | $0.694 \pm 0.061$ | 0.262 |
| 10 | $0.698 \pm 0.043$ | 0.268 |
| **15 (default)** | $\mathbf{0.712 \pm 0.028}$ | **0.296** |
| 20 | $0.706 \pm 0.039$ | 0.307 |
| 30 | $0.718 \pm 0.044$ | 0.311 |

**Correlation cutoff $\tau$ (with $K = 15$)**

| $\tau$ | AUC | Cross-fold Jaccard |
|---|---|---|
| 0.3 | $0.717 \pm 0.046$ | 0.247 |
| 0.4 | $0.703 \pm 0.034$ | 0.307 |
| **0.5 (default)** | $\mathbf{0.712 \pm 0.028}$ | **0.296** |
| 0.6 | $0.694 \pm 0.041$ | 0.288 |

Table S7: Robustness of GAE-Δ (shared-encoder default) to the choice of edge filter on the per-cancer effective universe (Methods, "Gene universe"): Pearson within-group correlation thresholding vs. Fisher's z differential-correlation filter (which corrects for both within-group variance and per-group sample size). All values are 10-fold CV AUC mean ± s.d. with $\tau = 0.5$.

| Edge filter | BLCA | BRCA | CESC | LIHC | LAML |
|---|---|---|---|---|---|
| Pearson (default) | $.703 \pm .061$ | $.719 \pm .046$ | $.726 \pm .067$ | $.718 \pm .039$ | $.722 \pm .070$ |
| Fisher's $z$ | $.695 \pm .074$ | $.713 \pm .055$ | $.717 \pm .080$ | $.710 \pm .051$ | $.700 \pm .085$ |
| Δ AUC | −0.008 | −0.006 | −0.009 | −0.008 | −0.022 |

Table S8: Multi-seed ensemble stability of GAE-$\Delta$ gene selection, computed under the shared-encoder default configuration on the per-cancer effective universe (Methods, "Gene universe"). For each cancer, GAE-$\Delta$ is run with five independent random seeds on the full cohort. "Inter-seed Jaccard" is the mean off-diagonal pairwise Jaccard of the per-seed top-100 selections; "5/5 seeds" is the number of genes selected in all five seeds; "$\geq$3/5 seeds" is the number of genes selected in at least three of the five seeds. "Cross-fold Jaccard (shared, single seed)" is reproduced from Table S11 (shared-encoder row) for direct comparison: inter-seed agreement is consistently $\approx 2\times$ cross-fold agreement, decomposing the total instability into a roughly equal data-driven and initialisation-driven component.

| Cancer | Inter-seed Jaccard | 5/5 seeds | $\geq$3/5 seeds | Cross-fold Jaccard (shared, single seed) |
|---|---|---|---|---|
| BLCA | $0.51 \pm 0.04$ | 43 / 100 | 87 / 100 | $0.211 \pm 0.04$ |
| BRCA | $0.54 \pm 0.03$ | 48 / 100 | 91 / 100 | $0.234 \pm 0.03$ |
| CESC | $0.49 \pm 0.04$ | 39 / 100 | 84 / 100 | $0.196 \pm 0.05$ |
| LIHC | $0.50 \pm 0.03$ | 41 / 100 | 88 / 100 | $0.202 \pm 0.04$ |
| LAML | $0.47 \pm 0.05$ | 36 / 100 | 79 / 100 | $0.183 \pm 0.05$ |

Table S9: GSEA prerank: top 10 MSigDB Hallmark pathways in LIHC ranked by FDR (IsoForest anomaly score across the 5 513 genes of the LIHC effective universe used as the natural background; see Methods, "Gene universe"). Nine of the ten reach FDR $q < 0.05$; Hypoxia ($q = 0.16$) is shown as the highest-ranked non-significant pathway for context. NES = normalised enrichment score; FDR $q$ = Benjamini–Hochberg-corrected $q$-value over 1 000 random-rank permutations.

| Hallmark Pathway | NES | FDR $q$ |
|---|---|---|
| IL-6/JAK/STAT3 Signaling | 2.77 | $1.0 \times 10^{-3}$ |
| Complement | 2.31 | $1.0 \times 10^{-3}$ |
| PI3K/AKT/mTOR Signaling | 2.23 | $3.0 \times 10^{-3}$ |
| Inflammatory Response | 2.16 | $4.4 \times 10^{-3}$ |
| Apoptosis | 2.04 | $4.4 \times 10^{-3}$ |
| Allograft Rejection | 2.08 | $5.3 \times 10^{-3}$ |
| Wnt-$\beta$-Catenin Signaling | 1.96 | $8.9 \times 10^{-3}$ |
| KRAS Signaling Up | 1.76 | 0.026 |
| Coagulation | 1.74 | 0.026 |
| Hypoxia | 1.43 | 0.16 |

Table S10: Number of MSigDB Hallmark pathways reaching FDR $q < 0.05$ per cancer in GAE-$\Delta$ GSEA prerank analysis (IsoForest anomaly score across each per-cancer effective universe; 1 000 permutations).

| Cancer | Hallmark FDR $< 0.05$ | Top pathway (NES) |
|---|---|---|
| BLCA | 7 | KRAS Signaling Up (2.41) |
| BRCA | 9 | Estrogen Response Early (2.62) |
| CESC | 6 | E2F Targets (2.31) |
| LIHC | 9 | IL-6/JAK/STAT3 Signaling (2.77) |
| LAML | 4 | Inflammatory Response (2.05) |

Table S11: Architectural ablation of the encoder-coupling and residual-correction stages of GAE-Δ across all five TCGA cohorts (10-fold CV). The default *Shared encoder, no KNN* configuration trains a single GAE jointly on both group graphs and reads out per-group embeddings from the same encoder, eliminating the inter-run alignment problem by construction. *Independent encoders* trains a separate GAE per phenotype group and is retained in the released codebase as a switchable option. "AUC mean / AUC median" are the mean and median across the 10 folds; the two columns track each other within $\leq 0.007$ AUC in all 15 cell pairs, indicating that no fold-level outlier drives the mean. The final column gives the one-sided paired Wilcoxon $p$-value (per-fold AUC) for the shared-encoder default > the row variant, and is left blank (−) for the default row itself. All paired tests fail to reach $p < 0.05$, indicating that the AUC differences between the three configurations are within fold-level noise; the shared-encoder default is preferred for architectural simplicity (no post-hoc alignment), and KNN correction is retained in the released codebase as an optional refinement that buys $\approx 0.10$ higher cross-fold gene-selection Jaccard at no significant AUC cost. The shared-encoder variant exhibits larger per-fold AUC variance (e.g. CESC $\sigma = 0.067$) than the KNN-corrected variant (e.g. CESC $\sigma = 0.028$); the rank-based Wilcoxon test that drives our conclusions is insensitive to this variance asymmetry, but readers prioritising low fold-to-fold AUC variance over architectural simplicity should prefer the independent + KNN variant.

| Cancer | Variant | AUC (mean) | AUC (med.) | Jaccard | $p_{\mathrm{w}}$ vs. shared default |
|---|---|---|---|---|---|
| BLCA | **Shared enc. (default)** | .703±.061 | .701 | .211 | – |
| | Indep. enc. + KNN | .697±.028 | .696 | .310 | .32 |
| | Indep. enc., no KNN | .694±.039 | .692 | .218 | .38 |
| BRCA | **Shared enc. (default)** | .719±.046 | .716 | .234 | – |
| | Indep. enc. + KNN | .714±.027 | .713 | .328 | .28 |
| | Indep. enc., no KNN | .708±.037 | .706 | .235 | .36 |
| CESC | **Shared enc. (default)** | .726±.067 | .720 | .196 | – |
| | Indep. enc. + KNN | .720±.028 | .719 | .301 | .31 |
| | Indep. enc., no KNN | .708±.040 | .706 | .214 | .41 |
| LIHC | **Shared enc. (default)** | .718±.039 | .716 | .202 | – |
| | Indep. enc. + KNN | .712±.028 | .711 | .304 | .25 |
| | Indep. enc., no KNN | .708±.041 | .704 | .212 | .39 |
| LAML | **Shared enc. (default)** | .722±.070 | .715 | .183 | – |
| | Indep. enc. + KNN | .716±.029 | .715 | .289 | .34 |
| | Indep. enc., no KNN | .712±.042 | .709 | .205 | .43 |

*For reference: LIHC, four-way comparison at $K = 15$, $\tau = 0.5$:*

| Variant | AUC | Cross-fold Jaccard |
|---|---|---|
| Shared encoder, no KNN (default) | $0.718 \pm 0.039$ | 0.202 |
| Independent encoders + KNN | $0.712 \pm 0.028$ | 0.304 |
| Independent encoders, no KNN | $0.708 \pm 0.041$ | 0.212 |
| Shared encoder + KNN | $0.694 \pm 0.035$ | 0.286 |

Table S12: Gene-set agreement between the default Pearson within-group correlation edge filter and the Fisher's $z$ differential-correlation filter on LIHC (effective universe; 5 513 genes). Consensus genes are those selected by Isolation Forest in at least 8 of 10 cross-validation folds. Driver hits and hypergeometric $p$-values are computed against the cancer-type-matched COSMIC CGC list intersected with the LIHC effective universe (the natural background, by construction). Jaccard is computed between the two consensus sets directly.

| Edge filter | Consensus size | Driver hits | Hypergeom. $p$ | Drivers found |
|---|---|---|---|---|
| Pearson (default) | 25 | 4 | $7.7 \times 10^{-5}$ | CDK1, MYC, CTNNB1, TP53 |
| Fisher's $z$ | 18 | 3 | $2.0 \times 10^{-3}$ | CDK1, MYC, TP53 |

Jaccard(Pearson, Fisher) = 0.40; shared drivers: CDK1, MYC, TP53.
Pearson-only driver: CTNNB1 (down-weighted by within-group variance correction).
